# Genomic and pedigree-based approaches to predict parental breeding values for nut and kernel traits in almond (*Prunus dulcis* Mill. D. A. Webb)

**DOI:** 10.64898/2026.01.22.701136

**Authors:** Shashi N. Goonetilleke, Melanie J. Wilkinson, Michelle G. Wirthensohn, Cassandra Collins, Agnelo Furtado, Robert J. Henry, Craig Hardner

## Abstract

The self-incompatibility, perennial growth habit, large tree size, and long juvenility present challenges in applying traditional breeding approaches in almond (*Prunus dulcis* Mill. D. A. Webb). Moreover, nut and kernel traits in almond are mainly controlled by a large number of small-effect quantitative trait loci (QTLs) and improving complex traits through conventional breeding approaches is slow and often inefficient. Genome-wide selection represents a promising strategy to enhance the efficiency of cultivar identification and selection of superior parents in almond breeding programs by estimating the breeding values (BVs) at early maturity. The main aim of this study was to implement genomic (GBLUP) and pedigree-based (ABLUP) prediction approaches to estimate BVs to identify the superior parental candidates for improving nut and kernel traits in almond. Here, we estimated BVs for nine traits that are commonly used in the primary evaluation stage of the almond breeding using genomic data from 61 parents and phenotypic data of 15,281 progeny derived from 205 unique families. Breeding values obtained from both approaches showed a strong correlation (*r* ≥ 0.94) for all traits except shell seal (*r* = 0.87). The population structure analysis conducted using high-quality 90K single nucleotide polymorphisms (SNPs) indicated clear separation of the Californian, European and some old Australian almond cultivars, with considerable admixture across some cultivars. Following further validation, both prediction approaches could be useful in early identification of superior candidates. The slightly higher breeding values obtained using the GBLUP compared to the ABLUP approach suggest that accounting for within-family variations and realised genomic relationships can enhance prediction accuracy, reliability, and overall genomic prediction performance in almond.

## INTRODUCTION

Almond (*Prunus dulcis* Mill. D. A. Webb), a member of the Rosaceae family, is the most widely consumed nut crop, with global consumption reaching 1.7 million tonnes in 2022 (Almond Board of California 2023). This popularity is attributed to its appealing colour, nutritional value, pleasant taste, versatility in culinary and confectionery applications. Between 2011 and 2023, global almond production increased by 54%, reaching 1.45 million metric tonnes of kernel (USDA 2024). Almond markets have different preferences, and the largest Asian almond markets prefer uniform size nuts covered with soft or paper shells while Mediterranean markets prefer almonds with hard shells (Almond Board of Australia 2023). In addition, the USA caters to customer-specific preferences with standard or unique sizes and shapes of nuts and kernels (Almond Board of California 2023). Therefore, improving kernel and nut quality traits is an important objective in almond breeding programs around the world.

To meet market preferences and consumer demands, almond breeding programs widely use approaches that involve controlled pollination between selected parents, evaluation of hybrid seedlings, and selection and 2_nd_ stage evaluation of vegetatively replicated superior clones prior to final selection for release (Wirthensohn and Sedgley 2002; Gradziel and Martínez-Gómez 2013). In almond breeding, phenotypic evaluation of nut and kernel traits cannot be undertaken until at least three years after planting due to the long juvenile period of almond (Goonetilleke and Fernández i Martí 2023). Selection based on molecular marker information allows earlier selection of individuals with desirable nut and kernel traits (Aranzana et al. 2019; Goonetilleke et al. 2023; Mas-Gómez et al. 2023). Currently, genetic improvement of almond is mainly done by identifying the associated genetic regions using quantitative trait locus (QTL) mapping approaches and/or association mapping approaches (Sánchez-Pérez et al. 2007; Font i Forcada et al. 2012; Goonetilleke et al. 2018; Sideli et al. 2023). Mapping of QTLs often captures larger effect loci but lacks the power to detect smaller effect loci and is often unable to provide an accurate picture of loci contributing to kernel and nut trait variation in crops (Xu et al. 2017). Most kernel traits seem to have polygenic inheritance with strong environmental effects (Goonetilleke et al. 2018; Sideli et al. 2023) that demand approaches that capture effects of all markers across the genome simultaneously for better understanding the genotype-environment effect and genetic improvement of these quality traits.

Estimated breeding values (EBVs) derived from pedigree (Henderson 1984) or genomic (Meuwissen et al. 2001; Kostick et al. 2023) information, represent a modern breeding approach that predicts the additive genetic merit for a trait that enables improved selection accuracy and accelerated genetic gain. In plant and animal breeding, EBVs using additive best linear unbiased prediction method (ABLUP) that derived from the pedigree-based relationship matrix and genomic-based best linear unbiased prediction (GBLUP) that uses genome-wide marker data (such as SNPs) to predict the can capture both major and minor loci effects for BV estimation are widely used. GBULP is used in genomic selection (GS) and can enable early selection of elite individuals for desired traits without phenotypic data, accelerates genetic gains and improve the efficiency of breeding program. When training the GBLUP models, phenotypic and genotypic information are jointly combined to estimate breeding values. GBLUP models assume markers are in close linkage with many QTLs of small effects that are random, independent and normally distributed. These models treat the collective effect of all markers as a single random effect, assuming equal variance across markers and normally distributed residuals. Linkage disequilibrium (LD) allows the markers to capture the effects of underlying QTLs. The genetic relatedness among individuals is calculated using a realised relationship matrix (GRM) derived from genotypic data. GBLUP leverages a genomic relationship matrix to model genetic similarity among individuals and provides more accurate assessment of genetic relationships. GBLUP also allows for more reliable distinction between full- and half-sibling relationships than ABULP as the GRM reflects the actual proportion of shared DNA to identify and manage hidden relationships such as inbreeding within populations and admixture that may not be directly identified by pedigree data alone (Meuwissen et al. 2001; Kostick et al. 2023).

Although several studies on marker-trait associations have been reported in almond using different types of populations, such as bi-parental crosses (Sánchez-Pérez et al. 2007; Wu et al. 2009; Tavassolian et al. 2010; Fernandez i Marti et al. 2012; Goonetilleke et al. 2018), and association panels (Font i. Forcada et al. 2015; Sideli, et al. 2023), to detect larger effect QTLs for subsequent selection of superior individuals with the improved kernel and nut traits, implementation of genomic prediction in almond is very limited. In this study, we applied ABLUP and GBLUP approaches to predict breeding values for a diverse panel of parents with known pedigree that that produced multiple phenotyped progeny. We compared the two approaches in terms of prediction accuracy and reliability and evaluated progeny performance for nine nut and kernel traits. Based on our findings, we propose strategies to improve early selection of superior individuals that facilitates shortening the breeding cycle across multiple generations and reduce the cost of field trials in almond.

## MATERIALS AND METHODS

### Study population

The F_1_ population used in the study consisted of 15,281 (Table S1) seedling progeny from 205 unique families derived from controlled crossing of 61 parents. The pedigree records (Tables S2-S3) were sourced from available literature and almond breeding programs in five countries: Australia, France, Italy, Spain and the USA. All the parents originated in other breeding programs were imported by the Australian Almond Breeding Program through collaborations.

The progeny and parents were established over multiple years from 1997 to 2018 in five randomised block breeding trial sites with some replication located at four locations in Australia: Dareton (34°04′57.7″S 142°00′55.0″E) in New South Wales, Lindsay Point (37°53′0″S 145°10′0″E) in Victoria, two sites in Monash (37°53′0″S 145°10′0″E) in Victoria and Loxton (34°27′14″S 140°34′01″E) in South Australia.

### Phenotypic data collection and analysis

Phenotypic data were collected from seedlings in four orchards from 1998 to 2019 from the trial sites were planted over multiple years and operated at different times over the 14-year assessment period. Therefore, continuous data were not available for all sites throughout the entire trial duration. Lindsay Point trial site was assessed from 1998 to 2001, Monash trial site was assessed from 2003-2007, Dareton trial site was assessed from 2007 to 2011, and Loxton trial site was assessed in 2019 (Table S3).

Each F_1_ progeny was assessed at least once for nine nut and kernel quality traits over the trial period: in-shell weight (ISW), shell weight (SW), percentage of double kernel (DK%), shell hardness (SH), kernel weight (KW), kernel taste (KT), testa colour (TC), surface appearance (SA), and shell seal (SS) of the kernel. For the assessment, 30 mature fruits were randomly collected from each tree to minimise sampling bias. The fruit was considered mature when the mesocarp was completely dried and split along the fruit suture and the peduncle was close to complete abscission. Samples were stored at room temperature for at least one month before assessment. All the assessments are carried out by the experienced one or two assessors. After weighting nuts using an electronic balance to obtain ISW, the nuts were cracked open to obtain KW and SW. For all these traits, the average value of the 30 nuts was used for the analysis. Percentage of double kernel (DK%) was obtained for each tree as a percentage by counting the number of shells containing the double kernels in a total of 30 nuts. Shell hardness (SH) was calculated as a percentage using the formula (kernel weight/nut weight) × 100%) proposed by Rugini (1986) and classified as follows: <25% (stone-hard), 25-35% (hard), 35-45% (semi-hard), 45-55% (soft), and <55% (paper shell), respectively.

Testa colour (TC) of the kernel was obtained by visually examining the depth of colour in the kernel and scored from 1 to 5 to indicate very light (5), light (4), medium (3), dark (2), and very dark (1), respectively with 0.5 increments. Kernel taste (KT) was assessed by chewing the kernel to taste and a score of 0 to 2 was given to indicate kernels with bitter (0), semi-bitter (1) and sweet (2), respectively with 0.5 increments. Shell seal (SS) was determined by visually examining the degree of shell closure that enclose the kernel using 1 to 5 scale with 0.5 increments: very good (5), good (4), better (3), bad (2) and very bad (1) seal, respectively. Surface appearance (SA) of the kernel was given a score from 1 to 10 based on kernel shape, smoothness, uniformity, pubescence, and overall visual acceptance.

### Genotypic data

#### Plant material, DNA extraction and genotyping

DNA from the 61 almond cultivars (Table S1) used as parents in the almond germplasm at the University of Adelaide, Waite Campus, Urrbrae, in South Australia (34°58′11″S 138°38′18″E) and Lindsay Point, Victoria (37°53′0″S 145°10′0″E) in Australia were extracted from young leaves using an in-house optimised CTAB protocol for yield and quality (Schultz et al. 2021). After DNA quality assessment by electrophoresis on 0.8% agarose (Bio-rad, USA) and measuring the 260/230 ratio using NanoDrop for DNA purity, and 36 cultivars were genotyped using whole genome resequencing to ∼20× coverage in NovaSeq 6000 sequencing platform at Ramaciotti Centre for genomics, Sydney, NSW, Australia, and the other whole genome resequences of 25 cultivars were obtained from a collaboration through Washington State University, USA and IRTA, Spain.

#### Genotypic data filtering and SNP calling

Before merging the pair-ended forward and reverse sequencing reads, the raw reads were trimmed by removing adapter sequences and low-quality sequences using BBMAP using the following parameters: Cleaned reads were then aligned to the reference genome Texas (pdulcis26.chromosomes.fasta) (Alioto et al. 2020) using BWA-MEM with default parameters v0.7.15-r1140 (Li and Durbin 2010). Mapped reads were then sorted, and duplicates were marked using Picard tools v2.17.0 (Broad Institute, 2017). Only reads that mapped to a unique position on the reference genome were retained for downstream analyses.

Identification of variants, including SNPs and indels, was performed using the Genome Analysis Toolkit (GATK) v.3.8 (McKenna et al. 2010) with the HaplotypeCaller method using the following quality control thresholds: minor allele frequency (MAF) <0.01, read depth <10, SNP heterozygosity <0.4, missing rate per SNP > 30% and monomorphic SNPs were removed. To minimize variant-calling errors caused by incorrect mapping and to extract original high-quality variants, genotypes of single-variant loci with read depth values less than 3 were removed using VCFtools v0.1.14 (Danecek et al. 2011) and base calls were recalibrated using 65,245 known SNPs from a previous almond GBS analysis (Goonetilleke et al. 2018).

#### Genotype imputation

Missing data were imputed using Beagle v4.1 (Browning and Browning 2016) with the following parameter settings: window = 15,000, overlap = 5,000, and iterations = 25. No pedigree information was used in the imputation procedure. Resulting dataset was tested for Hardy-Weinberg equilibrium (HWE) (*p*- value = 1e-6) using PLINK v1.90b6.24 (Purcell et al. 2007; Chang et al. 2015).

#### Population structure analysis

Principal component analysis (PCA) was performed to understand the population structure using PLINK v1.90b6.24 (Purcell et al. 2007; Chang et al. 2015) and R v4.1.3 (R Core Team, 2023) to calculate genetic distances between individuals based on identity-by-state and all first-degree relations were removed using the king-cutoff function. Population structure assessment was carried out using STRUCTURE v2.3.1 (Pritchard et al. 2000) by running 100 times on the SNP set with varying random seeds. CLUMPP v1.1.2 (Jakobsson and Rosenberg 2007) was used to align membership coefficients (Q-matrices) before clustering based on similarity. The final matrix of estimated admixture proportions was generated by averaging the matrices of the largest cluster.

#### Linkage disequilibrium

To identify redundant SNPs and independent haplotype blocks in the almond genome, linkage disequilibrium (LD) coefficients (r_2_) between pairwise high-quality SNPs were calculated using PopLDdecay v3.40 beta with default parameters (Zhang et al. 2019). The r_2_ was calculated for every combination of SNPs within a 250 kb window. LD decay graphs were produced using the Plot_MultiPop.pl program included in PopLDdecay v3.40.

#### The linear mixed-effects models

The general linear mixed effect model used to evaluate the similarity of pedigree and genomic based mixed linear models was:

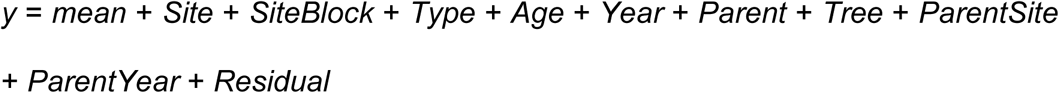

where *y* is the dependent trait, *Site* is the fixed site main effect across trees and years, *SiteBlock* is the fixed effect of the Block within Site, *Type* is the fixed main effect of the plant type (seedling progeny from different parent combinations or families), *Year* is the fixed main effect of year of assessment, *Parent* is the random mean additive genetic effect of parents of the progeny across sites and years (with distribution N (0, 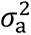)), *Tree* is the permanent non-additive genetic effect within sites and across assessment years (N (0, 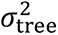)), *ParentSite* is the random additive genetic effect of a parent specific to a *Site* N (0, 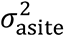)), *ParentYear* is the random additive genetic effect of a parent specific to an assessment year (N (0, 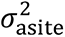)), and *Residual* is the common variance in *y* across sites and years not explained by the model (N (0, 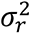).

The general form of the linear mixed model fitted to the progeny data to estimate genetic architecture and predict (pedigree or genomic based) breeding values was:

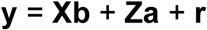

where **y** was a vector of the adjusted phenotypic data for each trait, **X** was a design matrix that maps the observations onto the unknown fixed effects (*mean*, *Site*, *SiteBlock* + *Type* + *Age* + *Year* ), **b** was a vector of the unknown fixed effects, **Z** was the design matrix for the random effects (*Parent*, *Tree*, *ParentSite*, *ParentYear*), **a** was the vector of the unknown random effects, and **r** was a vector of unknown random residuals.

The variance of **y** was:

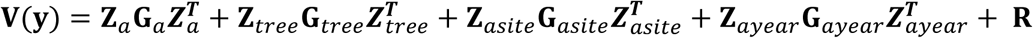

where **G***_a_* was the variance covariance among parental breeding values with the form

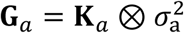

where **K***_a_* was the expected additive relationship matrix among parents estimated using the pedigree (**A**) or the realised additive genomic relationship matrix estimated following Van Raden (2008) (**GRM**), and 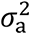 is the unknown genetic variance. The form of **G***_tree_* was

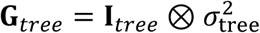

where **I***_tree_* was an identity matrix of dimensions number of trees,

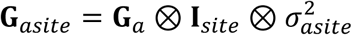

where **I***_site_* was an identity matrix of dimensions number of sites, and

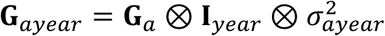

where **I***_year_* was an identity matrix of dimensions number of years.

The matrix A was constructed from the pedigree relationships using AGHmatrix v2.1.4 (Rampazo Amadeu et al. 2023) in R v4.2.1 (R Core Team 2023). To reduce collinearity and marker redundancy in the construction of the GRM from correlated markers, linkage disequilibrium weights were computed based on LD blocks identified using PLINK v1.90b6.24. SNP pruning was performed with a sliding window of 50 SNPs, a step size of 5 SNPs, and a marker correlation (r²) threshold of 0.2, retaining only SNPs in approximate linkage equilibrium. These pruned, largely independent SNPs were then used to construct a realised genomic relationship matrix (GRM) based on the VanRaden method (VanRaden 2008) with the ASRgenomics v1.1.0 package (Gezan et al. 2022) in R v4.2.1 (R Core Team 2023). Similarly, 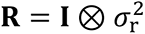. Unknown variance components were estimated using REML implemented in ASReml-R v4.2 (Butler et al. 2009) and used to predict pedigree or genomic based breeding values of the parents.

#### Preliminary analyses

For each site, preliminary analyses were undertaken using reduced linear models that omitted Site effects. To assess the assumption of normality underlying the F-statistic and to ensure unbiased predictions, the Shapiro–Wilk test was applied (Shapiro & Wilk 1965). Where the normality assumption was not supported, log transformations were used to improve residual alignment. Extreme residuals were investigated for abnormal experimental influences and rejected where this was established. To reduce the impact of heterogeneity of variance on genotype by year and genotype by site interactions, all phenotypic data were first scaled per site per year by dividing by site/year standard deviation (SD) (here after known as adjusted phenotypes).

#### Evaluating genetic effects

To examine how genetic effects are structured for each trait, two pedigree-based models were fitted. (i) to estimate among years, variance components were estimated using a year-specific pedigree model where data from each year were analysed separately to account for environmental variation among years (ii) to evaluate the genetic effects among sites, variance components were estimated using a site-specific pedigree model where data from each location were analysed separately to account for environmental variation among sites.

To evaluate the genetic effects using genomic models, data from each year were analysed separately to account for environmental variation among years. Similarly, data from each location were analysed separately to account for environmental variation among sites using a site-specific genomic model.

#### Narrow-sense heritability

To measure the proportion of phenotypic variance that is due to additive genetic variance that could reliably transmitted from parent to offspring, individual narrow-sense heritability (h_2_) for each of these traits were estimated.

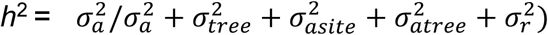

The genetic co-efficient of variation was measured for each trait as the additive genetic variance 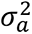 divided by the phenotypic mean across all the individuals. Narrow-sense heritability for both genomic and pedigree models were estimated using the pin function in R using ASReml-R v4.2 (Butler et al. 2009).

For all fitted models under both scenarios, the statistical significance of random effects was assessed using log-likelihood ratio tests. To evaluate the agreement between pedigree-based and genomic-based predictions, the estimated breeding values from both models were compared using Pearson correlation coefficient (*r*).

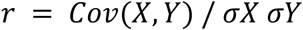

#### Prediction efficiency and unbiasedness

Prediction efficiencies of the ABLUP and GBLUP models were evaluated and compared based on the predictive ability, and predictive accuracy. Predictive ability was estimated by randomly partitioning the dataset so that either 50% or 75% of observations were used for model training, with the remainder used for validation. The two training proportions test the stability of model performance under different levels of data availability. Higher consistency across these splits indicates a more robust predictive model.

1. The predictive ability (*P1*) was defined as the Pearson correlation between the EBV of the individuals (EBVP) and their adjusted phenotypes (y). i.e.,

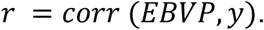

2. The predictive accuracy (*P2*) was estimated as the *r* scaled by h_2_ (square root of individual narrow sense heritability), i.e., *r*2/ = *corr*(*EBVP, y*)/√ℎ2 (Meuwissen et al. 2013).

In both pedigree and genomic approaches, unbiasedness of predictions was evaluated by regressing the observed values (true or adjusted phenotypes) on the estimated breeding values (EBVs). An intercept close to zero and a regression slope near one were used to indicate that the predictions were unbiased.

#### Implementing genomic prediction in almond breeding decision making process

To evaluate the effectiveness of incorporating the estimated breeding values into the almond breeding decision making process, we compared the number of progeny that were response to selection from the University of Adelaide (UoA) traditional selection procedure (C) with four other alternative selection strategies: (A) Nonpareil (commercial reference cultivar) + SD, (B) population mean + SD, (C) the UoA almond selection protocol, and (D) random selection at 50% cut-off point, and (E) 75% cut-off point (Table 7). The main aims were to identify individuals with high genetic potential and superior performance for each trait using EBVs and traditional procedures, and to evaluate the selection efficiency of both methods based on the number of parents selected (Table 7).

## RESULTS

### Phenotypic variation

Phenotypic values for the traits were highly consistent across years (*r* ≥ 0.95) (Figure S1). However, considerable phenotypic variation was detected among 15,281 progeny in a total of 205 unique families for all nine traits assessed (Tables S4-S5, Figure S1). Based on the measured phenotypic data, shell hardness phenotypes were positively correlated with kernel weight phenotypes but negatively correlated with shell weight phenotypes. Kernel weight phenotypes showed a strong negative correlation with in-shell weight (Table S6).

### Resequencing of the almond parents

Sixty-one almond parents generated 3.4 TB high quality reads with an average coverage of 15.8 reads per individual. Each almond cultivar generated an average of 4.7 GB data at approximately 20-fold depth. After conducting quality checks, the lowest sequence depth with raw data was obtained for Price (15.8×) and the highest was Constanti (23.8×). On average 99% of reads from each cultivar were mapped to the almond reference genome Texas. Overall, 9.8 million single-nucleotide polymorphisms (SNPs) were identified with an average of 41.67 SNPs per kilobase (kb) in the almond genome (Figure S2). These SNPs were distributed among all eight chromosomes, and each chromosome contained some regions with high-density SNPs of 750 or more within a region of 10 kb. After applying stringent hard filtering on the initial VCF, 125,580 SNPs resulted where coverage ranged from 13× (Mandaline and R1065) to 39× (Gaura). Removing of multiallelic and highly co-related SNPs, through LD pruning, a total of 75,596 high-quality, independent SNPs with a minor allele frequency ≥ 0.05 were identified for the downstream analysis. SNP validation using the known SNP sites and the Texas v.1 genome assembly revealed an accuracy of 96.9%. Following genotype imputation, the accuracy of heterozygous SNPs was markedly improved, and the SNP-calling accuracy was increased to 98.8%.

### Population structure and linkage disequilibrium

Haploblock analyses identified a total of 6,806 with an average block size of 7.79 kb with the longest haploblock being 163.83 kb. The linkage disequilibrium (LD) analysis indicated rapid LD decay in the almond genome about 300 kb (*r*_2_ = 0.158) and the mean pairwise *r*_2_ values ranged from 0.145 to 0.172 across all chromosomes (Figure S3). LD decay varied across the eight chromosomes, being most rapid on Chromosome 2 and slowest on Chromosome 6 (Figure S4).

The first two principal components of the PCA, explained 25.05% of the variation (Figure 1). The old Australian almond cultivars (Chellaston, Somerton, Parkinson, Johnston’s Prolific, Mckinlays) were clearly separated from the rest of the almond cultivars, and the new almond cultivars originated from the crossing of the European and USA parental materials placed in between two major clusters of the European and USA almond cultivars. Among the different clusters obtained, the European cluster that contained almond cultivars from Spain, Italy and France showed high genetic similarities and cultivars from different European breeding programs did not separate clearly. The Narrowest genetic diversity within cluster was detected in USA cluster.

**Figure 1:**
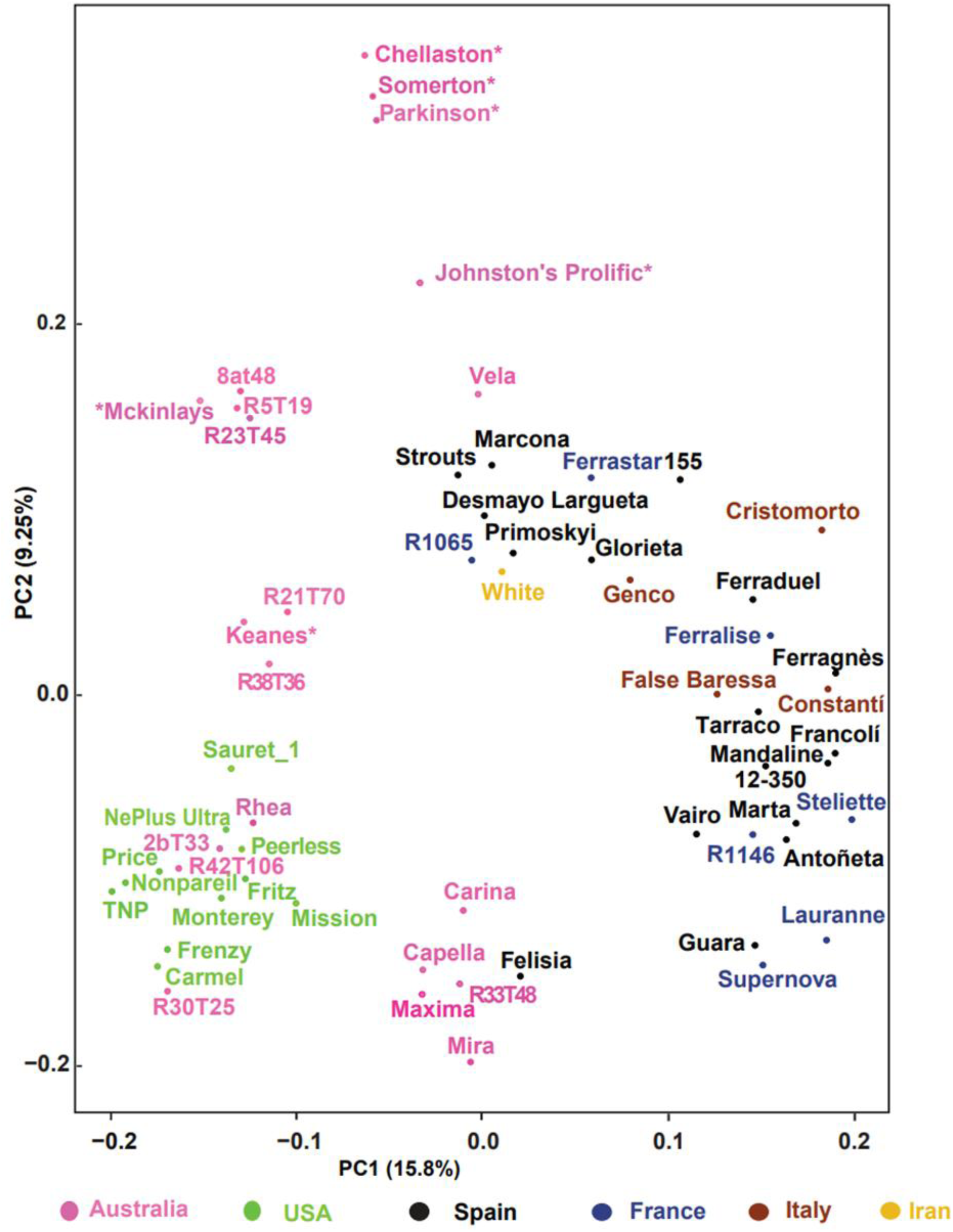
Principal component analysis of the almond clones/cultivars used in the study using over 80K single nucleotide polymorphism markers. Almond clones/cultivars from Australia, USA, Spain, France, Italy and Iran are shown in magenta, green, black, blue, brown and beige, respectively. Old Australian almond cultivars that were selected based on best performing phenotypes are indicated by an asterisk.

The kinship coefficient matrix (not shown) resulted from the Structure (K = 4 clusters) indicated a wide range of values from 0.08 to 0.98, with an average 0.48. As expected, the genetic composition of some of the almond cultivars revealed considerable admixture pattern across some investigated cultivars (Figure 2). Furthermore, the diagonal and off-diagonal values of the A matrix were 1 and ranged from 0 to 0.5, respectively, whereas the GRM diagonal and off-diagonal values ranged from 0.98 to 1.5 and -0.05 to 0.52, respectively.

**Figure 2:**
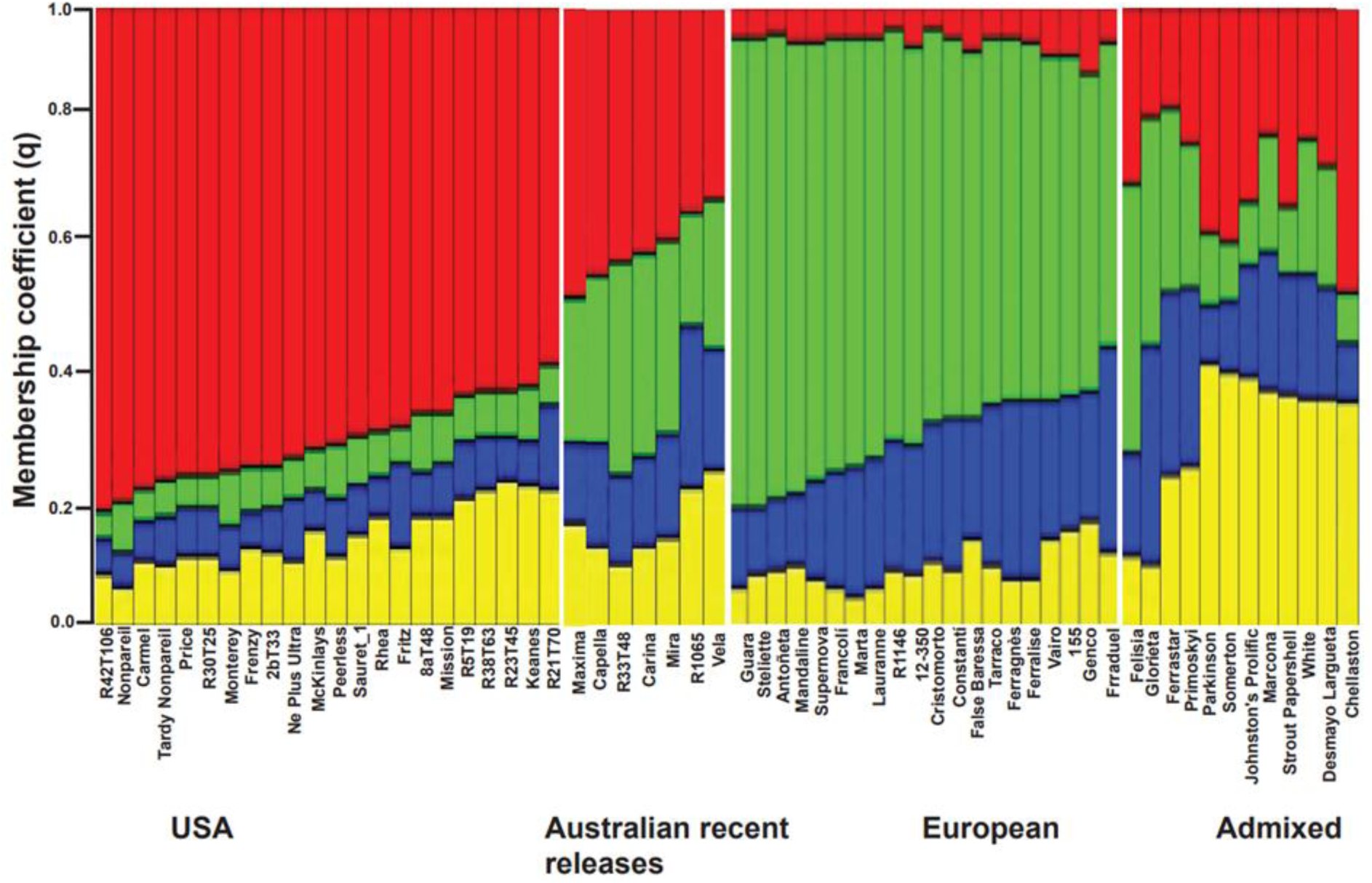
Population structure analysis for almond clones/cultivars used in the study using over 80K single nucleotide polymorphism (SNP) markers (K = 4). Each cultivar is represented by a vertical bar. The length and the colour of segments in each bar represent the proportion of the genome (q) contributed by each of four ancestral populations. Cultivars assigned to one of the populations have a membership coefficient (q) for that population >0.6. The remaining cultivars are classified as admixed.

### Explained phenotypic variation in two approaches

The histogram and Q-Q plots (Figure S5) for quantitative traits indicated that random effects were normally distributed. Furthermore, despite differences in the inferred genetic architecture of these nut and kernel traits, approximate normality of the estimated random effects detected using histograms and Q-Q plots support the suitability of the mixed-model assumptions.

### Genetic architecture of nut and kernel traits

In both approaches, the proportion of phenotypic variance explained by the genetic effects was similar with the models that have site × year as fixed terms, (Figure 3, Table 3). In both approaches, the highest proportion of phenotypic variance were obtained for shell weight (54%), whereas the lowest was detected for the surface appearance, 24% for genome-based approach and 28% for pedigree-based approach, respectively. In both approaches, except in-shell weight, kernel weight, shell weight and testa colour, the residual component observed were higher than the genetic component (genotype) and had the highest impact on overall variation observed i.e. from 43% (in-shell weight and kernel weight) to 66% (shell hardness) in pedigree model and 74% (surface appearance) to 44% (kernel weight) (Table 3).

**Figure 3:**
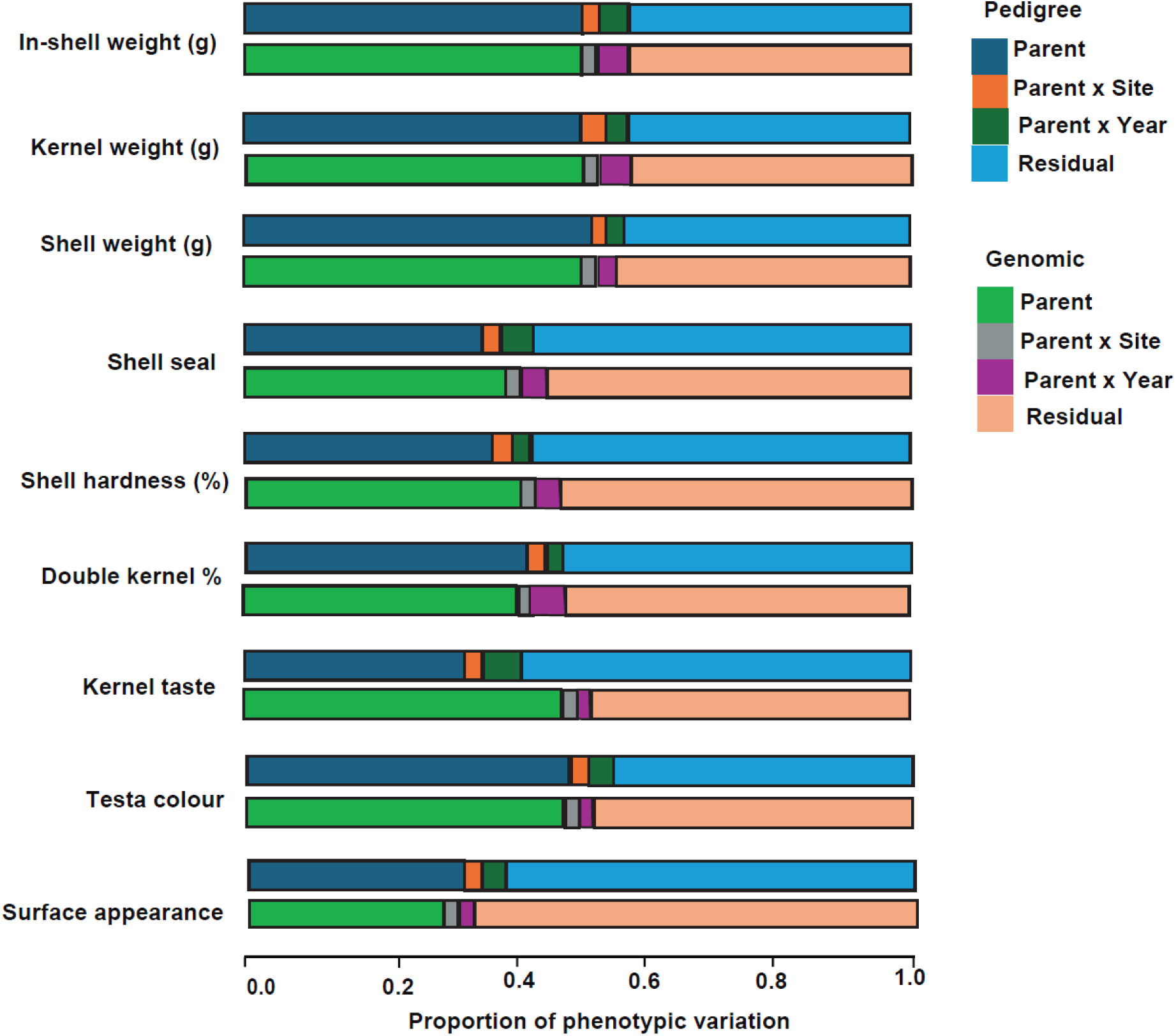
Proportion of phenotypic variation explained by the components in the pedigree-based (upper panel) and genomic based (lower panel) full models for each trait.

The deviation from the average genotypic effect across all years by the genotypic effect within years accounted for 0.3% (in-shell weight) to 1.1% (shell harness) in pedigree-based model and 0.01% (in-shell weight) to 1.3% (shell hardness) in genomic-based model. With site effects and individual family within year effects were both significantly below 9%. In the pedigree-based method this component ranged from 0.001% (testa color) to 8.3% (surface appearance) whereas, in the genomic-based models these values varied from 0.01% (shell seal) to 1.5% (surface appearance). Confirming the phenotypic variation, the highest mean narrow-sense heritability of 0.54 was detected for in-shell weight using all models (Tables 2) and the lowest of 0.27 for surface appearance of the kernel (Table 2). There were strong negative correlations between predicted breeding values of kernel weight and in-shell weight (*r*= -0.95, p = 0.000001) (Figure 4). There was also a non-significant negative correlation between the predicted breeding values for shell hardness and shell weight (*r*= -0.37, p = 0.21).

**Figure 4:**
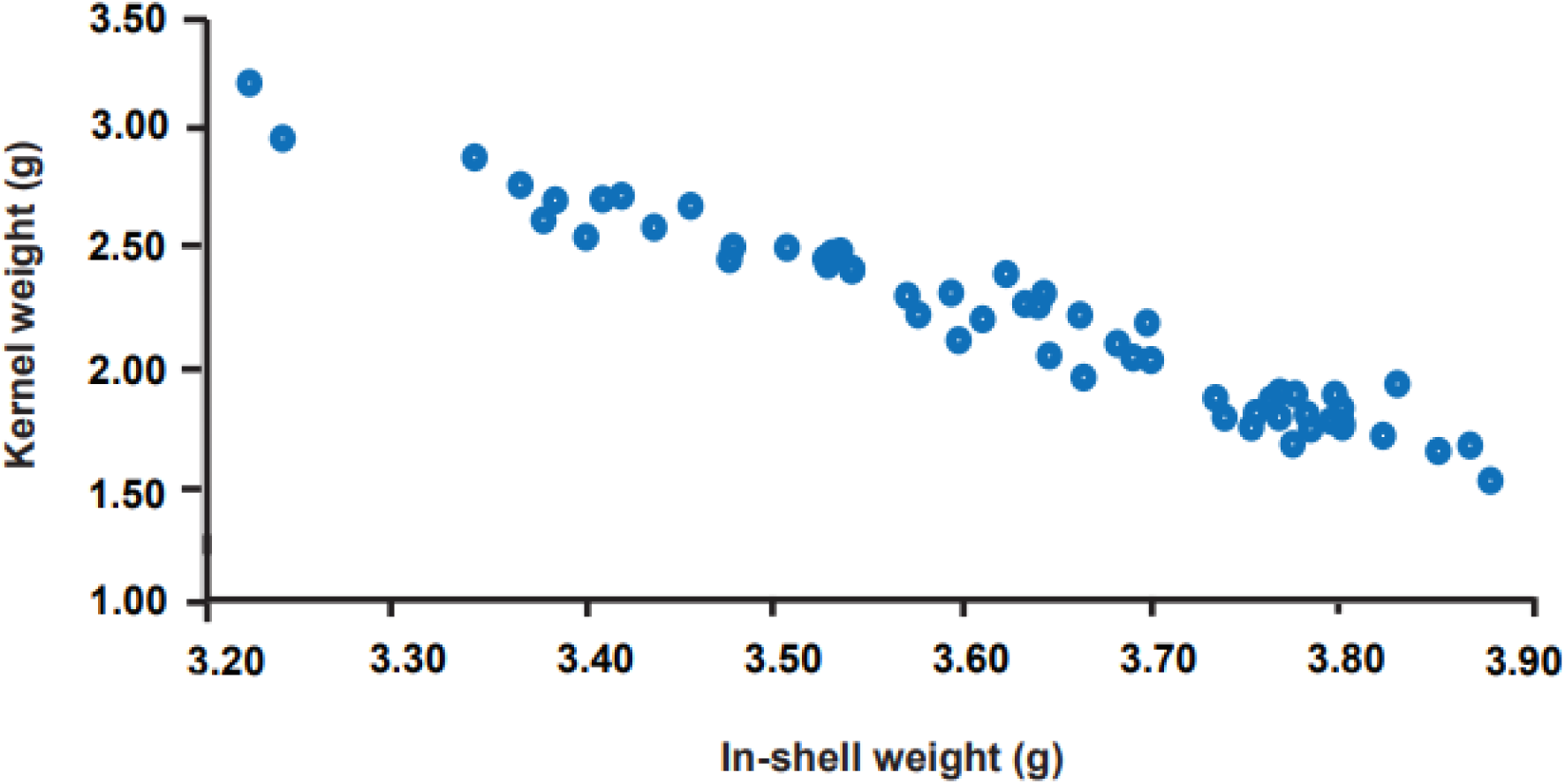
Strong negative correlation observed between estimated breeding values of in-shell weight and kernel weight (*r*=-0.95).

**Table 1.** Estimations of narrow-sense heritability (*h*^2^), additive variance, residual variance and log-likelihood ratio test (LRT) obtained as a measure of the overall model fit for the pedigree-based models. For *h*^2^ Standard error (SE) is given.

| Trait | $h^2$ | | | Additive variance | | | Residual variance | | | Loglikelihood ratio | | |
| --- | --- | --- | --- | --- | --- | --- | --- | --- | --- | --- | --- | --- |
|  | Full model | Year specific | Site specific | Full model | Year specific | Site specific | Full model | Year specific | Site specific | Full model | Year specific | Site specific |
| In-shell weight | 0.52±0.01 | 0.54±0.01 | 0.48±0.01 | 16850.52 | 16992.93 | 16902.09 | 21758.32 | 21165.63 | 21911.53 | -15211.78 | -15212.58 | -15212.69 |
| Kernel weight | 0.51±0.01 | 0.50±0.08 | 0.49±0.01 | 16751.21 | 17281.35 | 16952.25 | 23363.12 | 20765.32 | 23126.32 | -15210.78 | -15210.78 | -15211.78 |
| Shell weight | 0.34±0.11 | 0.32±0.02 | 0.34±0.11 | 16950.52 | 17184.91 | 16962.11 | 21798.43 | 21465.59 | 21912.57 | -15210.92 | -15210.25 | -15211.79 |
| Shell seal | 0.41±0.12 | 0.35±0.11 | 0.36±0.11 | 15652.31 | 15861.42 | 15668.23 | 25816.32 | 26116.03 | 26916.93 | -15211.85 | -15214.34 | -15213.79 |
| Shell hardness | 0.37±0.12 | 0.36±0.26 | 0.35±0.12 | 17963.45 | 17245.25 | 17243.45 | 26245.31 | 26198.32 | 26564.35 | -15010.32 | -14980.78 | -15011.65 |
| Double kernel % | 0.44±0.11 | 0.47±0.26 | 0.45±0.11 | 15932.42 | 15982.35 | 15981.25 | 22145.98 | 22178.39 | 22264.47 | -15012.31 | -15231.45 | -15231.21 |
| Kernel taste | 0.37±0.11 | 0.33±0.11 | 0.35±0.12 | 15958.56 | 15758.41 | 15832.51 | 26123.78 | 26071.38 | 26125.58 | -15012.31 | -15221.45 | -15222.38 |
| Testa colour | 0.52±0.09 | 0.54±0.09 | 0.53±0.09 | 15421.68 | 15425.28 | 15481.59 | 22752.53 | 21278.41 | 22138.52 | -15202.31 | -15201.45 | -15202.39 |
| Surface appearance | 0.28±0.09 | 0.27±0.09 | 0.29±0.09 | 15489.41 | 15981.25 | 15681.45 | 28925.78 | 28578.39 | 28235.41 | -15206.28 | -15205.45 | -15206.12 |

**Table 2.** Estimations of narrow-sense heritability (*h*^2^), additive variance, residual variance and log-likelihood ratio test (LRT) obtained as a measure of the models it for the genomic-based models. For *h*^2^ Standard error (SE) is given.

| **Trait** | **$h^2$** |  |  | **Additive variance** |  |  | **Residual variance** |  |  | **Log-likelihood ratio** |  |  |
| --- | --- | --- | --- | --- | --- | --- | --- | --- | --- | --- | --- | --- |
| Model | Full model | Year specific | Site specific | Full model | Year specific | Site specific | Full model | Year specific | Site specific | Full model | Year specific | Site fixed |
| In-shell weight | 0.48±0.01 | 0.51±0.06 | 0.46±0.02 | 17850.52 | 17938.11 | 17859.12 | 23417.2 | 21473.85 | 23543.1 | -14236.21 | -14236.66 | -14236.18 |
| Kernel weight | 0.50±0.07 | 0.54±0.07 | 0.52±0.02 | 16936.25 | 17532.6 | 17126.32 | 22187.35 | 22237.25 | 22235.68 | -14238.27 | -14237.61 | -14238.39 |
| Shell weight | 0.52±0.08 | 0.54±0.08 | 0.55±0.08 | 15526.32 | 15529.92 | 15527.32 | 22789.32 | 21878.85 | 22335.21 | -14239.10 | -14237.65 | -14238.43 |
| Shell seal | 0.39±0.52 | 0.36±0.15 | 0.38±0.12 | 15623.51 | 15663.25 | 15662.31 | 27912.34 | 25256.1 | 26861.3 | -14238.23 | -14127.63 | -14127.81 |
| Shell hardness | 0.41±0.15 | 0.40±0.15 | 0.41±0.15 | 16865.45 | 17287.58 | 17293.42 | 23236.63 | 21497.32 | 23562.35 | -14120.23 | -14037.61 | -14038.39 |
| Double kernel % | 0.39±0.08 | 0.39±0.15 | 0.39±0.09 | 16022.58 | 16182.35 | 16280.78 | 23045.44 | 24024.41 | 23164.58 | -14238.01 | -14137.60 | -14235.31 |
| Kernel taste | 0.32±0.09 | 0.35±0.09 | 0.34±0.11 | 15940.41 | 16082.55 | 16065.25 | 24145.98 | 24579.51 | 24264.37 | -14138.20 | -14037.61 | -14039.33 |
| Testa colour | 0.50±0.09 | 0.49±0.09 | 0.48±0.09 | 16032.42 | 16423.34 | 16232.41 | 26254.92 | 24458.29 | 25061.85 | -14215.91 | -14217.63 | -14215.31 |
| Surface appearance | 0.24±0.09 | 0.25±0.09 | 0.25±0.11 | 15802.58 | 15985.68 | 15879.25 | 26443.91 | 27088.39 | 27164.57 | -14395.25 | -14373.65 | -14381.85 |

### Correlation of predicted breeding values

Both ABLUP and GBLUP resulted in similar estimated breeding values (Figure S6) with strong positive correlations. The Pearson correlation coefficient (*r*) varied from 0.87 (shell seal) to 0.98 (shell hardness) (Table S7, Figure S7).

### Prediction abilities and accuracies in two approaches

Predictive abilities derived from the random subsets of pedigree-based and genomic-based full models showed variation across the evaluated almond quality traits (Table 4). The GBLUP model showed slightly higher predictive abilities than the ABLUP model for most traits, with an average increase of 0.03. For kernel weight, predictive ability improved from 0.42 (ABLUP) to 0.56 (GBLUP), corresponding to an accuracy gain from 0.60 to 0.71. Similarly, GBLUP improved predictions in-shell weight, shell weight, shell hardness, double kernel, and kernel trait while ABLUP slightly outperformed GBLUP for testa colour and surface appearance (Table 4).

**Table 3.** Variance (random and residual) components estimated as a percentage using pedigree-based and genomic-based full models fitted for each trait.

| **Trait** | **Model** | **Component** |  |  |  |
| --- | --- | --- | --- | --- | --- |
|  |  | **Parent** | **Parent × Site** | **Parent × Year** | **Residual** |
| In-shell weight | Pedigree-based | 51.37 | 1.16 | 3.71 | 43.74 |
| Kernel weight | Pedigree-based | 51.11 | 1.57 | 3.52 | 43.78 |
| Shell weight | Pedigree-based | 54.12 | 0.04 | 1.71 | 44.12 |
| Shell seal | Pedigree-based | 35.71 | 0.05 | 0.62 | 63.62 |
| Shell hardness (%) | Pedigree-based | 32.21 | 0.88 | 0.89 | 66.03 |
| Double kernel % | Pedigree-based | 47.90 | 0.15 | 0.15 | 51.79 |
| Kernel taste | Pedigree-based | 33.45 | 1.78 | 0.28 | 64.49 |
| Testa colour | Pedigree-based | 53.94 | 1.16 | 0.01 | 44.90 |
| Surface appearance | Pedigree-based | 28.18 | 0.52 | 8.33 | 62.96 |
| In-shell weight | Genomic-based | 51.12 | 0.03 | 3.26 | 45.18 |
| Kernel weight | Genomic-based | 51.01 | 0.59 | 3.87 | 44.53 |
| Shell weight | Genomic-based | 53.57 | 0.01 | 1.01 | 45.41 |
| Shell seal | Genomic-based | 39.40 | 0.10 | 0.11 | 62.37 |
| Shell hardness | Genomic-based | 41.90 | 1.29 | 1.28 | 55.52 |
| Double kernel % | Genomic-based | 39.47 | 0.82 | 1.50 | 58.19 |
| Kernel taste | Genomic-based | 32.54 | 0.38 | 0.43 | 66.63 |
| Testa colour | Genomic-based | 50.02 | 0.573 | 1.12 | 48.23 |
| Surface appearance | Genomic-based | 24.12 | 0.22 | 1.53 | 74.12 |

**Table 4.**
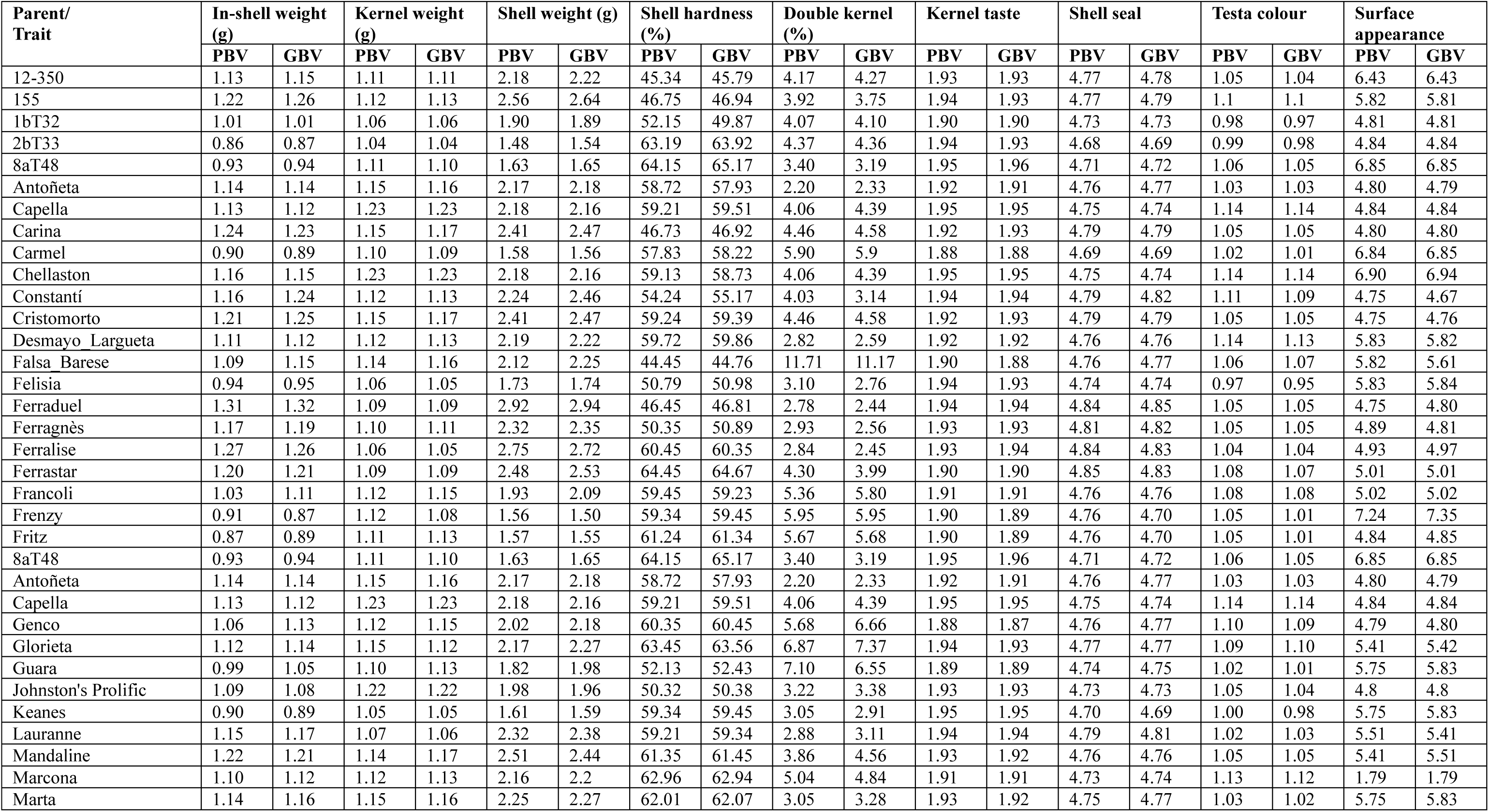
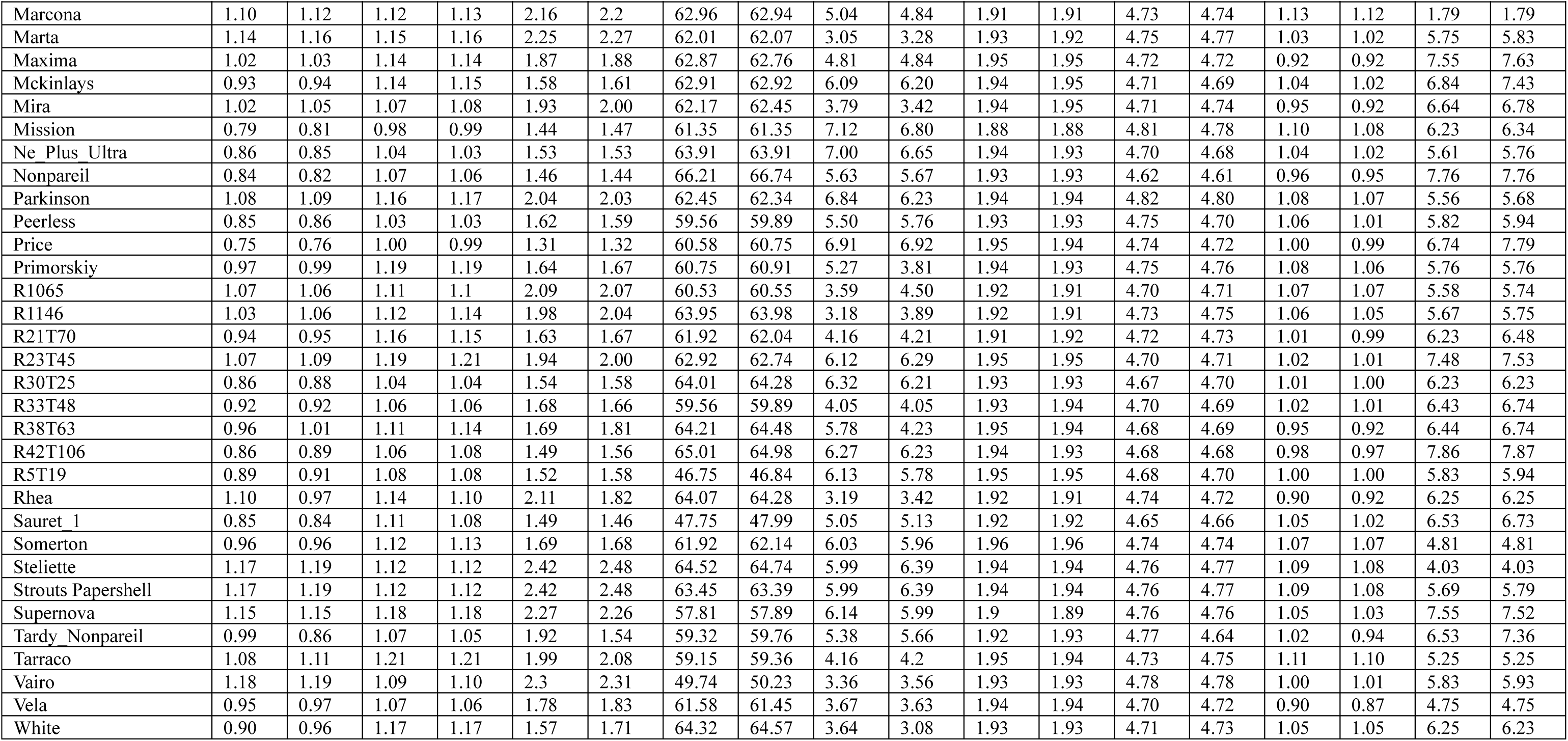
Estimated Breeding values obtained using pedigree-based (PBV) and genomic-based (GBV) approaches using full model.

Overall, genomic prediction achieved moderate to high accuracies (0.4-0.7), confirming its utility for improving almond quality traits. When 50% of the population with both genotypic and phenotypic data was used as the training set and the validation set included only genotypic data, the mean predictive abilities ranged from 0.35 to 0.58. Increasing the training set proportion to 75% resulted in higher mean predictive abilities, ranging from 0.44 to 0.65 (Table 4). Further, when the complete genotypic dataset (n= 75,596 SNPs) was available, GBLUP produced 30-38% accuracy over nine traits. When the size of the validation population increased (50% to 75%), the accuracy obtained also increased (Table 5). Both pedigree-based and genomic-based models produced nearly unbiased predictions, with regression slopes close to unity (0.85-0.97). Minor over-dispersion was observed for surface appearance, suggesting a slight overestimation of genetic differences for this trait (Table 6).

**Table 5.** Summary of predictive abilities (*r*) of pedigree and genomic based full models using randomly selected validating set for almond kernel and nut traits.

| Trait | Validating set scheme | | Predictive ability ($r$) | | | |
| --- | --- | --- | --- | --- | --- | --- |
|  | Method | Validating set | Mean | SD | Minimum | Maximum |
| In-shell weight (g) | Random selection | 50% | 0.46 | 0.03 | 0.34 | 0.58 |
|  |  | 75% | 0.56 | 0.05 | 0.34 | 0.70 |
| Kernel weight (g) | Random selection | 50% | 0.42 | 0.03 | 0.34 | 0.50 |
|  |  | 75% | 0.45 | 0.05 | 0.34 | 0.62 |
| Shell weight (g) | Random selection | 50% | 0.53 | 0.03 | 0.44 | 0.61 |
|  |  | 75% | 0.56 | 0.04 | 0.40 | 0.66 |
| Shell seal | Random selection | 50% | 0.41 | 0.03 | 0.32 | 0.50 |
|  |  | 75% | 0.44 | 0.05 | 0.32 | 0.59 |
| Shell hardness (%) | Random selection | 50% | 0.57 | 0.03 | 0.42 | 0.69 |
|  |  | 75% | 0.58 | 0.04 | 0.45 | 0.71 |
| Double kernel (%) | Random selection | 50% | 0.49 | 0.03 | 0.39 | 0.59 |
|  |  | 75% | 0.51 | 0.05 | 0.37 | 0.62 |
| Kernel taste | Random selection | 50% | 0.49 | 0.03 | 0.40 | 0.62 |
|  |  | 75% | 0.52 | 0.04 | 0.42 | 0.69 |
| Testa colour | Random selection | 50% | 0.49 | 0.03 | 0.37 | 0.56 |
|  |  | 75% | 0.51 | 0.05 | 0.36 | 0.67 |
| Surface appearance | Random selection | 50% | 0.47 | 0.03 | 0.35 | 0.56 |
|  |  | 75% | 0.51 | 0.05 | 0.36 | 0.67 |

**Table 6.** Summary of predictive accuracy (*r/*√*h*^2^) of pedigree and genomic based full models using randomly selected validating set for almond kernel and nut traits

| Trait | Validating set scheme | | Predictive accuracy ($r/\sqrt{h^2}$) | | | |
| --- | --- | --- | --- | --- | --- | --- |
|  | Method | Validating set | Mean | SD | Minimum | Maximum |
| In-shell weight (g) | Random selection | 50% | 0.58 | 0.04 | 0.39 | 0.71 |
|  |  | 75% | 0.57 | 0.05 | 0.39 | 0.71 |
| Kernel weight (g) | Random selection | 50% | 0.56 | 0.03 | 0.39 | 0.69 |
|  |  | 75% | 0.57 | 0.05 | 0.39 | 0.71 |
| Shell weight (g) | Random selection | 50% | 0.53 | 0.03 | 0.44 | 0.73 |
|  |  | 75% | 0.56 | 0.04 | 0.40 | 0.72 |
| Shell seal | Random selection | 50% | 0.51 | 0.03 | 0.41 | 0.67 |
|  |  | 75% | 0.44 | 0.05 | 0.39 | 0.65 |
| Shell hardness (%) | Random selection | 50% | 0.53 | 0.03 | 0.44 | 0.61 |
|  |  | 75% | 0.56 | 0.04 | 0.40 | 0.66 |
| Double kernel (%) | Random selection | 50% | 0.48 | 0.03 | 0.37 | 0.56 |
|  |  | 75% | 0.52 | 0.05 | 0.36 | 0.67 |
| Kernel taste | Random selection | 50% | 0.49 | 0.03 | 0.41 | 0.57 |
|  |  | 75% | 0.52 | 0.04 | 0.40 | 0.64 |
| Testa colour | Random selection | 50% | 0.48 | 0.03 | 0.37 | 0.56 |
|  |  | 75% | 0.52 | 0.05 | 0.36 | 0.67 |
| Surface appearance | Random selection | 50% | 0.44 | 0.03 | 0.37 | 0.51 |
|  |  | 75% | 0.46 | 0.05 | 0.36 | 0.54 |

**Table 7.** Unbiasedness evaluated by regressing observed phenotypes on estimated breeding values from genomic and pedigree based full models.

| **Trait** | **Model** | **Regression slope** |
| --- | --- | --- |
| In-shell weight (g) | Genomic | 0.97 |
|  | Pedigree | 0.91 |
| Kernel weight (g) | Genomic | 0.93 |
|  | Pedigree | 0.92 |
| Shell weight (g) | Genomic | 0.96 |
|  | Pedigree | 0.94 |
| Shell seal | Genomic | 0.96 |
|  | Pedigree | 0.94 |
| Shell hardness (%) | Genomic | 0.96 |
|  | Pedigree | 0.92 |
| Double kernel (%) | Genomic | 0.97 |
|  | Pedigree | 0.91 |
| Kernel taste | Genomic | 0.92 |
|  | Pedigree | 0.91 |
| Testa colour | Genomic | 0.92 |
|  | Pedigree | 0.89 |
| Surface appearance | Genomic | 0.85 |
|  | Pedigree | 0.91 |

**Table 8.** Breeding goals and selection threshold applied in decision-making process for almond quality traits.

| Trait | Breeding goal | Selection threshold |  |  |  |  |
| --- | --- | --- | --- | --- | --- | --- |
|  |  | A | B | C* | D | E |
| In-shell weight | large with less shell weight | Nonpareil +SD | population mean +SD | UOA almond selection threshold | Top 50% selected | Top 75% selected |
| Kernel weight | large kernel | Nonpareil +SD | population mean +SD | UOA almond selection threshold | Top 50% selected | Top 75% selected |
| Shell weight | medium weight to obtain more kernel weight | Nonpareil +SD | population mean +SD | UOA almond selection threshold | Top 50% selected | Top 75% selected |
| Shell seal | Fully covered shell seal | Nonpareil -SD | population mean -SD | UOA almond selection threshold | Top 50% selected | Top 75% selected |
| Shell hardness (%) | Various | Keep all >25% |  |  |  |  |
| Double kernel % | Low <5% | Nonpareil -SD | population mean -SD | UOA almond selection threshold | Top 50% selected | Top 75% selected |
| Kernel taste | Sweet | Nonpareil +SD | population mean +SD | UOA almond selection threshold | Top 50% selected | Top 75% selected |
| Testa colour | Uniform light | Nonpareil +SD | population mean +SD | UOA almond selection threshold | Top 50% selected | Top 75% selected |
| Surface appearance | Very good overall visual and tactile characteristics | Nonpareil +SD | population mean +SD | UOA almond selection threshold | Top 50% selected | Top 75% selected |

C\* :University of Adelaide almond breeding primary evaluation selection threshold

### Genomic prediction in almond breeding decision making process

Genomic prediction could be employed to select high performing clones/cultivars for five important traits in the Australian almond breeding program with high confidence (Tables 4-6). Examination of genomic and pedigree-based estimated breeding values (Figure S6), along with the corresponding expected genetic gain or loss plots in the clones/cultivars (Figures S8, Table S8), indicated that this population can be selected for kernel weight, in-shell weight, shell weight, and shell hardness with high confidence using genomic prediction. It would partially reduce the tedious phenotyping efforts in the primary and secondary stages of the almond breeding (Figures S9-S10). Thus, deploying genomic prediction could reduce the length of the almond breeding cycle by at least 6 years considering that female flowering starts at 3 years from planting (Figures S9 and S10).

The number of parents and families selected or rejected depended on the selection method (conventional phenotypic vs. genomic selection; Table S9). Fewer individuals were rejected based on genomic predicted values than on conventional phenotypic trait values (Tables S11 and S12). Nevertheless, regardless of the selection method, the decisions to select or reject parents and families were largely consistent, occurring for approximately 45 to 85% and 55 to 87%, respectively.

## DISCUSSION

Here, we estimated breeding values for nine important nut and kernel traits that are used in the primary evaluation stage in almond breeding using both genomic and pedigree-based methods. This study is the to estimate breeding values using genomic and/or pedigree-based methods for the nut and kernel traits in almond. The population of 15,281 almond clones was ideal for estimating the reliable breeding values in this study, as it was large, genetically diverse, well-phenotyped, and well-connected, with accurate pedigrees and dense genomic information. It was from four different regions in Australia and structured to capture genetic variation while minimising bias from environmental effects or inbreeding. The similar EBVs obtained from the ABLUP and GBLUP methods indicate the potential of implementing prediction approaches in almond breeding, thereby enhancing the efficiency of the almond breeding decision-making process.

### Whole genome resequencing and population structure

The implementation of different sequencing platforms has been reported for almond such as genotyping-by-sequencing (GBS) (Goonetilleke et al. 2018; Pavan, et al. 2021), whole genome re-sequencing (Castanera et al. 2024) and SNP array (Duval et al. 2023). Here, we used 80K SNPs obtained from whole genome resequencing of 61 cultivars for genomic prediction of nut and kernel quality traits in almond. Other than the whole genome re-sequencing of 81 almond cultivars reported by Castanera et al. (2024), our study represents the largest resequencing efforts in almond clones with different origins.

In accordance with the results obtained from the pedigree information by Pérez de los Cobos et al. (2021) and Pavan et al. (2021) the clusters obtained from the principal component analysis indicated almond has at least three main origins: California (USA), Europe and Australia. Most of the recent Australian cultivars have founders from the USA and Europe, where most of the old Australian varieties such as Parkinson, Chellaston, Somerton, and Johnston’s Prolific are admixed. The clustering of European almond cultivars from Spain, France, and Italy was consistent with previous findings, reflecting their derivation from the same founding parents during the establishment of various almond breeding programs in Europe (Pavan et al. 2021; Pérez de los Cobos et al. 2021).

Consistent with Khojand et al (2024) and Pavan et al. (2021), using Iran and European almond populations, respectively, linkage disequilibrium (LD) analysis revealed a rapid decay of LD, with an average decay of ∼300 kbp at *r*^2^ = 0.158, highlighting the cross-pollinating nature of almond. Compared with related self-pollinated crop species, almond exhibits more rapid LD decay due to less constrained recombination. LD decay differs across chromosomes and Chromosome 2 had a very rapid decay and Chromosome 6 had the slowest decay, indicating variation in the recombination rate between chromosomes. The slowest decay on Chromosome 6 could be due to balancing selection for the *S*-locus that controls self-incompatibility in almond (Khojand, et al. 2024). Slow LD decay at the *S*-locus helps maintain allelic diversity and locus functionality by preserving regions of reduced recombination throughout evolution (Goonetilleke et al. 2023).

The off-diagonal values of the A matrix from pedigree-based relationships (0.5 for full-sibs, 0.25 for half-sibs) provides stable, expectation-based values that do not depend on environment or dataset that derived entirely from recorded pedigree, and assuming Mendelian inheritance. In contrast, GRM from genomic relationships that based on realised genomic relationships can be varied more, depending on location or data type, due to the segregation, recombination, and marker effects.

### Pedigree and genomic prediction in almond

The pedigree-based approach provided an efficient framework for estimating breeding values of nut and kernel quality traits, and its strong correlation with genomic-based estimates underlines the robustness of the genomic prediction. Estimated breeding values from these two methods showed a high level of correlation, indicating the suitability of these models in estimating the breeding values for future predictions. However, the main limitation in this study was that the clones used to train the genomic model were the same as those used for estimating breeding values in the pedigree-based approach and due to this overlap, the correlation between genomic based estimations and pedigree-based breeding values might appear higher than it would in an independent population. The genomic model may not be able to generalise to completely independent populations. Furthermore, estimated breeding values were obtained for each trait separately, considering only additive genetic variation as the genetic component. Because each EBV reflects the genetic merit of a clone for a single trait in isolation, combining these values into a unified selection index or to support a common breeding objective can be challenging. This is when traits are genetically correlated and/or breeders aim to improve multiple traits simultaneously, since simple aggregation of individual EBVs may not adequately capture trade-offs or the overall genetic potential of a clone across all target traits.

In developing performance prediction models, phenotyping primarily serves to train the prediction models, and the accuracy of phenotypic assessment is critical for predicting reliable breeding values and achieving long-term genetic gains. In our study, certain traits (kernel taste, shell seal, overall appearance, testa colour) were visually assessed and recorded as categorical variables. Because visual scoring can be subjective and may emphasise certain observable classes over others, this introduces ascertainment bias, which can potentially reduce the accuracy of genomic predictions. Using next generation high-throughput phenotyping (HTP) techniques for assessing these traits could be a more reliable approach. High-throughput phenotyping (HTP) technologies, including automated imaging systems, sensor-based measurements, and robotic platforms, enable rapid and efficient collection of phenotypic data Based on research conducted in cattle using high-throughput phenotyping (HTP) methods, prediction accuracy increased by 30-40% compared with models using traditional categorical phenotypes (Kizilkaya et al. 2014).

Generally, genomic prediction models assume that traits are continuous and approximately normally distributed. Deviations from this assumption, such as categorical or skewed traits, can reduce the accuracy of predicted breeding values. Linear mixed models offer the advantage of accounting for population relatedness, thereby reducing false-positive associations. However, practically, in the real-world not all data is normally distributed nor act independently. Genetically, all these traits are correlated, and many are controlled by the same loci (Fernandez i Marti et al. 2012; Goonetilleke and Fernández i Martí 2023). Therefore, it is important to investigate all genetic factors and potential nonlinear effects such as gene networks, regulatory mechanisms, and climatic conditions that can influence phenotypic expression, including epistatic and dominance interactions. Since, many of the almond nut and kernel traits are genetically correlated, selection based on the correlated response which accounts for the effect of selection on one trait on other genetically correlated traits represents an optimal strategy for identifying superior parents for breeding. Further, a multivariate analysis approach, which considers more than one trait simultaneously when fitting mixed models, could also be an effective strategy for achieving genetic gain across multiple traits at the same time.

### Predictive abilities for nut and kernel quality traits

The regression slopes obtained were close to unity for most almond quality traits, ranging from 0.85 to 0.97, indicating that both the pedigree-based and genomic-based full models produced mostly unbiased predictions. Minor deviations from one were observed for surface appearance, suggesting slight under- or over-dispersion of the predicted values for these traits. Overall, the genomic-based model (GBLUP) showed improved predictive ability and maintained unbiasedness comparable to the pedigree-based model, confirming its reliability for predicting genetic merit across almond quality traits.

For most nut and kernel quality traits, moderate predictive abilities were observed across randomly selected validation sets indicating that genome-wide selection can be an effective breeding approach for certain traits. In addition, all almond nut traits studied here were affected by a large number of moderate to small effect QTLs (Fernandez i Marti et al. 2012; Goonetilleke and Fernández i Martí 2023). Although the population used in this study was large, genetically diverse, and well-connected, most clones and families were unreplicated. Consequently, the observed variation among families reflected both genetic and environmental components. In addition, the predictive ability for each trait may have been influenced by factors such as the germplasm used, trait distribution, heritability, experimental design, and the statistical models applied. While direct references for comparison with our study were not identified in previous publications, the predictive abilities obtained in this study were comparable to values reported for similar traits in related crops such as apple (Roth, et al. 2020; Jung, et al. 2022). For example, the mean predictive abilities obtained for testa colour in this study ranged from 0.48 to 0.52 (Table 4), which were similar to the predictive abilities observed for skin colour reported in apple. i.e. Minamikawa, et al. (2024) reported predictive ability 0.57 for apple skin colour by considering only the additive genetic effects, across the 15 families in the apple breeding program in Japan (Minamikawa, et al. 2024). The testa colour variation in almond is primarily determined by the regulation of anthocyanin biosynthesis (Petroni and Tonelli 2011; Yan, et al. 2021). Among the key regulators, basic helix-loop-helix (bHLH) and WD40 transcription factors form part of the MYB-bHLH-WD40 (MBW) complex, which controls the expression of structural genes involved in anthocyanin synthesis. Similar regulatory mechanisms influencing fruit or seed coat colour have been reported in apple (*Malus domestica*) (McClure et al. 2019) and peanut (*Arachis hypogaea*) (Ahmad et al. 2023) and reported to have moderate heritability.

As expected, the large residual variance observed for all traits was not fully accounted for by the combined fixed effects of year and site on the genotypes (families). This suggests that the models could be further improved, for example, by incorporating additional genetic or environmental effects. The unexplained variance factors could be due to the differences caused by the location, within and between field effects, variations in management practices, and weather conditions. As these trials were conducted in different locations in Australia: Lindsay Point and Monash (VIC), Dareton (NSW) and Loxton (SA) from 1998 to 2019. However, these trials were not replicated. Conducting multi-environment (MET) and multi-location trials (MLT) would allow estimation of within- and between-location, as well as field-level, variation. Including these factors would improve the accuracy of genomic prediction models.

Over the trial period, this study was conducted across four locations in Australia with very similar weather conditions. For most traits, high main effects, and low genotype × environment (G×E) interactions indicated that genotypes behaved consistently across environments. However, the trials were not replicated and including MET and multi-location trials (MLT) would allow estimation of within and between location, as well as field-level variation, providing more accurate models for genomic prediction. Incorporating phenotypic data from multiple trials and international almond breeding programs would further capture environmental diversity and improve our understanding of the genetic determinants of traits. This approach also helps mitigate potential confounding effects from residual variance, as observed in this study. Including G×E interactions in the models is important, as predicted breeding values can be adjusted if significant G×E effects are detected.

Incorporating additional phenotypic data from multiple sources provides an opportunity to better understand G×E interactions. Evaluating individual performance across different locations allows more generalisable breeding value predictions and provides insight into trait stability across environments. (Budhlakoti et al. 2022; Hardner et al. 2022). Given high correlations obtained from the pedigree and genomic based methods for nine nut and kernel traits, genomic models can be applied widely. However, the adaptability and accuracy of these models need to be validated using almonds with diverse genetic backgrounds, as most of the clones used were derived from genetically similar parents, which may limit their applicability to progeny with higher genetic diversity.

### Implementing genomic selection in almond breeding decision making

Almond breeders simultaneously target multiple traits when making selection decisions in primary and secondary evaluation stages. The University of Adelaide almond breeding program uses their own threshold for each trait and overall ranking for primary stage selection of individuals. Therefore, predicting which individuals would provide overall high-quality kernel is challenging and not all traits can be targeted with genome-wide selection. However, considering the agreements in explained phenotypic variations from estimated breeding values with the values obtained from the assessments of phenotypic data, moderate narrow-sense heritability and the high correlation obtained with the pedigree-based and genomic-based approaches, genomic selection could be used to select some traits (such as kernel weight, in shell weight, shell weight and shell hardness) in the primary and secondary stages of the almond breeding program.

Generally, new almond clones or cultivars are selected against the benchmark cultivar Nonpareil without additional stringent thresholds. However, regardless of the selection method, the choice of thresholds significantly affected the number of individuals advanced for further evaluation. In this study, using trait levels of Nonpareil and the population mean resulted in the rejection of fewer parents and families compared to applying a 75_th_ percentile threshold for kernel weight and in-shell weight, which led to a 56% rejection rate. When Nonpareil trait levels were used in genomic selection (GS), the rejection rate was further reduced to 8%.

Further, candidate parents and proposed breeding program using EBVs from this study can be used to reduce the breeding cycle of almond and by surpassing the extensive phenotypic evaluation of each stage, the proposed breeding program has the potential of reducing the current conventional breeding cycle of 20 years to 14 years. One limitation of this selection study was the assumption that all traits carry equal breeding importance. In reality, the relative importance of each trait depends on the breeding objective and its impact in a given context. Further research is therefore needed to define trait-specific breeding targets and selection thresholds, including for multiple-trait scenarios, to facilitate the adoption of genomic selection in almond breeding decision-making.

This study highlighted that genomic prediction could provide an effective approach to enhance breeding efforts by leveraging genetic information to predict phenotypic traits in parents and families in the Australian almond breeding program. Although further validation is required, by integrating genomic data with traditional breeding methods, breeders and researchers can expedite the selection of desirable traits in almond efficiently. Furthermore, genomic selection can be used for early selection of seedlings by predicting breeding values before phenotypes are fully expressed. However, it requires careful model design with larger and more diverse training populations, multi-location, multi-environment data, and dense markers, along with proper validation. Machine learning can improve genomic prediction in almond by capturing nonlinear relationships and interactions among SNPs. Using methods such as random forests or deep learning with genomic and phenotypic data can increase the accuracy of breeding value predictions compared to traditional linear models. Moreover, genomic prediction could greatly enhance the efficiency of crop improvement efforts in tree crop species and the methods applied in this study can be easily adapted to other tree crop species.

## DATA AVAILABILITY

All data generated or analysed during this study are included in the published article and its supplementary information.

## ACKNOWLEDGEMENTS

Authors thank Jana Kolesik, University of Adelaide almond breeding program for assisting with phenotypic data collection and Hort Innovation (the grower-owned, not-for-profit research and development corporation for Australian horticulture) for financial support through the almond industry and development levy and contributions from the Australian Government and the University of Queensland.

## FUNDING

This project has been funded by Hort Innovation, using the tree genomics (LP_27032025) research and development levies, contributions from the Australian Government and co-investment from the University of Queensland. Hort Innovation is the grower-owned, not-for-profit research and development corporation for Australian horticulture.

## CONFLICT OF INTEREST

The authors declare no competing interests.

## AUTHOR CONTRIBUTIONS

C.H. conceived and designed the study, M.G.W. and S.N.G. developed the populations. S.N.G. and M.J.W. analysed the sequence data, S.N.G. and C.H. undertook the statistical analysis, C.H. and R.J.H. obtained the funding for sequencing, A.F. supervised the resequencing of almond and S.N.G wrote the first draft of the manuscript. All authors read and revised the manuscript.

